# Proliferation drives quorum sensing of microbial products in human macrophage populations

**DOI:** 10.1101/2022.05.12.491598

**Authors:** Nadia Rajab, Linden J. Gearing, Ruqian Lyu, Yair D.J. Prawer, Paul W. Angel, Sean M. Grimmond, Andrew L. Laslett, Davis J. McCarthy, Christine A. Wells

**Affiliations:** Centre for Stem Cell Systems, Department of Anatomy and Physiology, Faculty of Medicine, Dentistry and Health Sciences, The University of Melbourne, VIC, 3010, Australia; Centre for Innate Immunity and Infectious Diseases, Hudson Institute of Medical Research, Clayton, VIC, 3168, Australia; Department of Molecular and Translational Sciences, Monash University, Clayton, VIC, 3168, Australia; Bioinformatics and Cellular Genomics, St Vincent’s Institute of Medical Research, Fitzroy, Australia; Melbourne Integrative Genomics, School of Biosciences – School of Mathematics & Statistics, Faculty of Science, University of Melbourne, Melbourne, Australia; University of Melbourne Centre for Cancer Research, Faculty of Medicine, Dentistry and Health Sciences, The University of Melbourne, VIC, 3010, Australia; Australian Regenerative Medicine Institute, Monash University, Melbourne, VIC 3800, Australia

**Keywords:** Macrophage, pluripotent stem cells, endotoxin tolerance, innate memory, inflammation, lipopolysaccharide

## Abstract

Macrophages coordinate the initial host inflammatory response to tissue infection, as well as mediating the reparative phase, by producing growth factors that promote tissue repair. One model of this functional dichotomy is that peripherally recruited monocyte-derived macrophages drive acute inflammatory responses to infection, whereas tissue-resident macrophages are responsible for tissue repair. Alternatively, inflammation and repair may be inter-dependent molecular programs, such that both recruited and resident cells have equivalent capacity to contribute. Repeated exposure to pathogenic challenge results in innate tolerance, which may also alter the contributions of discrete macrophage populations to inflammation or repair. In this study a village model of tissue resident and recruited macrophages was created using induced pluripotent stem cell-derived macrophages and peripheral blood monocyte-derived macrophages, respectively. Population responses to repeated exposure to lipopolysaccharide were assessed with single-cell RNA sequencing and donors demultiplexed with Vireo. A subset of genes escaped classical tolerance programs in the iPSC, but not monocyte-derived macrophages, and this was associated with differences in their proliferative capacity. This suggests that targeting the proliferative resident macrophages would be most effective to limit inflammatory signaling.

## Introduction

Multiple tissue-resident macrophage subsets have been described and defined by roles that are specialized to their tissue of residency, such as bone resorption by osteoclasts^1^, neural pruning by microglia^2^, lipid-associated macrophages^3^, stem cell licensing in bone marrow^4^ and intestine^5^, or iron recycling in splenic macrophages^6^. Differences between macrophage subsets have also been described in the context of ontogeny (reviewed in ^7^), anatomical location, or molecular phenotype^8^. Given the diverse roles of macrophages in tissue development, homeostasis, and in host defense, there is considerable interest in understanding mechanisms that might target macrophage subsets in tissue inflammation or repair.

Functional differences between tissue-resident macrophages (TRM) and monocyte-derived macrophages (MDM) recruited in response to inflammation are controversial. In some models, such as brain, the resident macrophages (microglia) play a neuroprotective role, while recruited MDM are thought to be the inflammatory mediator^9^. In contrast, others have described diverse monocyte and macrophage subsets associated with age-related inflammation, cancer or trauma^10^, as well as inflammatory phenotypes associated with differences in IRF8 signaling across these subsets^11^. If macrophages have predefined roles in infection or inflammation, then this observed heterogeneity might well be explained by mixing of ontogenically and phenotypically distinct populations, exemplified by the progression of liver disease^12^. Alternatively, if both resident and recruited macrophage populations are intrinsically heterogeneous, then quorum-sensing will lead to roles that are randomly assigned across all populations, ensuring that the capacity for inflammation and repair is available regardless of the origin of the responding macrophage^13^.

This idea of quorum sensing is supported by early studies of stochasticity of macrophage responsiveness showing heterogeneity in gene expression after exposure to lipopolysaccharide (LPS). Examples of this have been described in studies showing variability of LPS responses in clones of murine RAW264.7 cells^13,14^. In the first study, subclones demonstrated varying levels of gene expression after LPS stimulation, suggesting the coexistence of macrophage subsets that could be propagated as subpopulations. In the second study, heterogeneity in the timing of RelA recruitment to the TNF promoter in a RAW264.7 reporter line was observed after LPS stimulation, but variability between cells could also be explained by population density. These studies demonstrate that multiple mechanisms impact on the timing, duration, and amplitude of macrophage activation, even in simple models receiving a highly polarizing inflammatory signal, such as LPS.

It is obvious that, in a tissue setting, macrophages will be integrating signals from the local environment and it is difficult to deconvolute the influence of these signals from ontogeny over the course of an inflammatory event. Tissue culture models offer an opportunity to address cell-intrinsic sources of macrophage diversity. Macrophages derived from human induced pluripotent stem cells (PSCM) present new opportunities to model tissue residency and factors that shape human macrophage development and activation. PSCMs are an excellent model of tissue residency because they share molecular ontogeny with TRM ^15,16^ and are functionally equivalent to TRM in engraftment models in brain and lung^17^. A recent comparison of macrophage transcriptomes in our own Myeloid Atlas confirmed that PSCMs shared the transcriptional phenotype of primary TRM including tissues such as brain, joint and liver^18^.

Given PSCMs offer an *in vitro* representation of tissue-resident cells, they provide a relevant experimental model to investigate sources of population heterogeneity in bacterial responses. While others^19^ have shown that PSCMs recognize and respond to LPS, it is not known whether they can be tolerized on repeated exposure, which would more accurately mimic the regulatory program seen after an infection than a single high dose pathogen exposure. We were also curious to see how coordinated acute responses to LPS were across human macrophage populations, as perhaps early versus late responders within a population could explain some of the previously described heterogeneity in trained monocytes^20^. Here, we use single cell RNA-sequencing to investigate population responses to high-dose LPS activation and then re-stimulation using a coculture, or village, model of PSCMs and MDMs, as models of tissue residency versus recruited macrophages respectively.

## Results

### Establishing a ‘village’ of macrophages to assess macrophage heterogeneity at the single cell level

The transcriptional heterogeneity of macrophage subsets was evident even in the unstimulated populations (Figure 1). The use of a village model removed population density as a possible driver of heterogeneity (Figure 1A), as all the cells were exposed to the same local environment, maintained in CSF-1 (C, control), or exposed to LPS for 2 hours (A, acute) or 18 hours (R, rest), Tolerised (T) macrophages were acutely stimulates as per (A and R), then washed after 18hours, and rested for 2 hours then restimulated for 2 hours with LPS. PSCM and MDM formed distinct clusters across the time course that were not explained by fractions of undifferentiated or nonresponsive cells – MDM and PSCMs share similar phenotypes of CD14+ CD16low populations after several days culture in CSF1 (Figure 1B). Cells collected at each time point were barcoded for demultiplexing after single cell RNA-sequencing and donor origin of the cell was determined by SNP deconvolution, to confirm that the heterogeneity was not driven by donor (Supplementary Figure 1A). LPS exposure was a strongly polarizing signal in the acute time periods, but population heterogeneity increased in the resting and restimulated conditions (Figure 1C, D, E).

**Figure 1.**
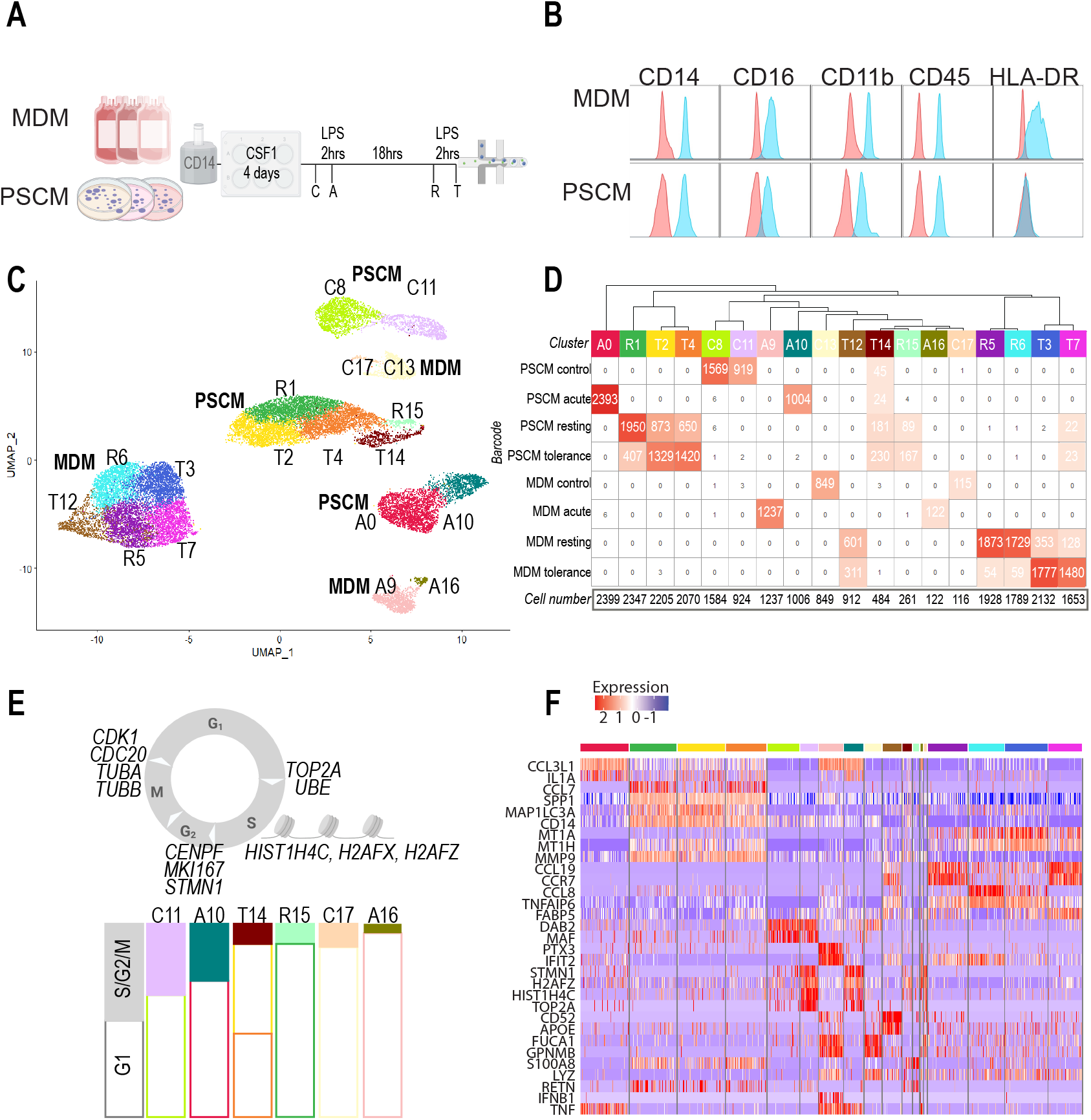
Proliferation drives macrophage heterogeneity in a village-in-a-dish model of repeated bacterial stimulation. (1A) Schematic of experimental design, including design of village coculture model for 3 MDM donors or 3 PSCM lines. Time course for LPS stimulation and multiplexed single cell library preparation. (1B) Representative surface expression of macrophage markers in MDM and PSCM measured by flow cytometry. Isotype control antibody staining (pink), signal (blue). (1C) Uniform Manifold Approximation and Projection (UMAP) of Seurat clusters (resolution 0.9) (1D) Seurat cluster group (columns) by sample barcode (rows). Groups are colour-coded according to (C) and labelled according to time point (C control, A acute, R 18 hours, T tolerance) and numbered from highest to lowest cell number per group. (1E) Genes that discriminated proliferating-cell clusters across LPS stimulation time course. Stacked bar plots indicate the proportion of proliferating-cell clusters (shaded bars) per barcode group. (1F) Key discriminating markers between UMAP clusters. Heat map of top genes (avg. Log2FC) in each subcluster. Legend denotes high (red) to low (blue) expression. See also Supplementary Figure 1, Supplementary Tables 1 and 2.

PSCM expressed hallmark genes of tissue resident macrophages (Figure 1F, Supplementary Table 1), including transcription factors associated with terminal macrophage differentiation, such as *MAF* ^21^, and markers of perivascular and lymphatic-associated macrophages including *CD169/ SIGLEC1, LYVE1* and *FOLR2* ^22,23^. PSCMs were transcriptionally similar to MDM and primary TRM (Supplementary Figure 1B and C) using the benchmarking tool in the Stemformatics Myeloid Atlas. MDM subsets were distinguished by expression of immunoregulatory glycoproteins *GPNMB*, a negative regulator of TLR signaling ^24^, macrophage migratory factors such as *FUCA1*, as well as chemokine receptors. However, the most striking difference between MDM and PSCM clusters was the presence of a proliferative subset of PSCM in the control, acute and tolerized conditions (Figure 1E).

The observations of ~40% of PSCM and <10% of MDM proliferating in the control condition is consistent with the CSF1 responsiveness of tissue resident vs recruited macrophages. PSCM and MDM were cultured in the presence of recombinant CSF1, which is a proliferative signal to tissue-resident macrophages and essential for the maintenance of bone marrow and circulating monocytes (reviewed in ^25^). Monocytes are postmitotic cells in circulation, although a small fraction have been identified proliferative in response to CSF1^26^. Just 10% of the PSCM were proliferating by the resting or tolerized conditions (Figure 1E), consistent with a pro-survival but antiproliferative consequence of TLR signaling. Both proliferating and nonproliferating PSCM subsets expressed equivalent levels of components of the TLR4-LPS pathway, at similar levels to the MDM control subsets (Figure 2A).

**Figure 2.**
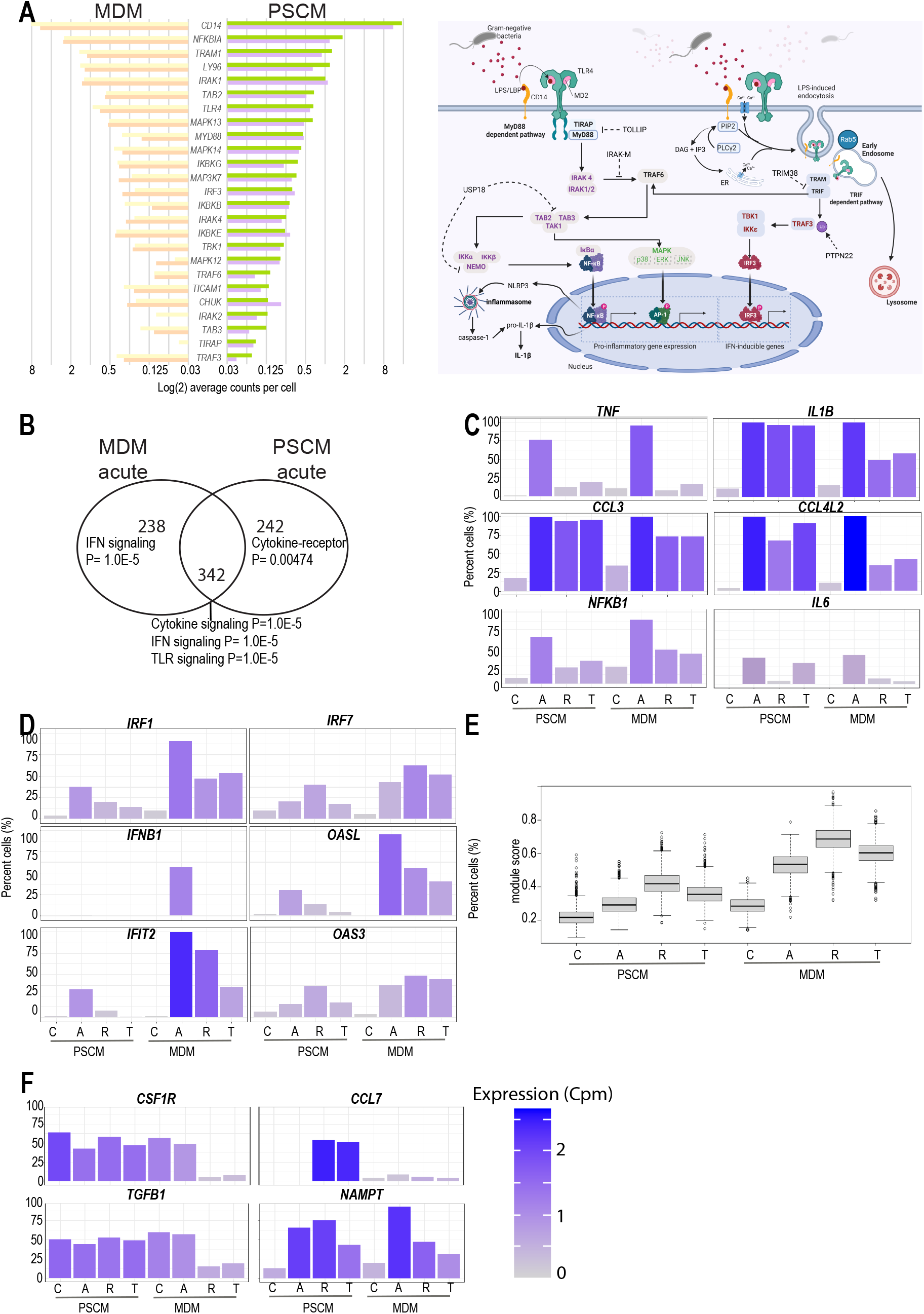
Inflammatory gene expression varied by the proportion of cells recruited to the inflammatory phenotype, as well as average gene expression per cell. (2A) Schematic of the TLR pathway and average expression of pathway members in unstimulated PSCM and MDM populations. Bar plots coloured by cluster members (MDM Cluster C13, C17; PSCM clusters C8, C11: see Figure 1C) to evaluate average expression in proliferating and nonproliferating control clusters. (2B) Venn diagram of LPS-inducible genes after 2 hours (selected by Log2FC. > 1.5, Adj. p value <0.05). P-values on graph indicate InnateDB pathway over-representation (Fishers exact test). (2C) Percentage of cells per cluster expressing six representative targets of TLR4-LPS stimulation. (2D) Percentage of cells per cluster expressing six representative targets of IRF (MyD88-independent) stimulation. (2E) Seurat module scores of the Reactome Interferon signaling pathway. Averaged cell score shown. (2F) Growth factor signaling persists in PSCM but not MDM. Percentage of cells per cluster expressing four representative reparative factors. For all bar plots, gene expression (average CPM / cell) indicated by colour where dark blue indicates highest expression, light for lowest expression.

### Proliferating PSCM escape LPS tolerance

LPS is a synchronizing signal that results in high expression of core inflammatory mediators in all cells receiving the signal. These included *TNF, IL1B, IL6, CXCL1, CXCL2, CXCL8, CXCL10, CCL3, CCL4L2*, and members of the shared interleukin/TLR signaling pathway including *NFKBIA* and *NFKB1* (Figure 2, Supplementary Table 1), and negative regulators of cytokine signaling such as *SOCS1* and *SOCS3*. These are predominantly targets of the TLR-MyD88-NF-κB pathway, demonstrating that the PSCM expressed an active TLR4 signaling cascade. After 18 hours activation, most inflammatory mediators have reached a baseline expression level in both macrophage populations. Restimulation with LPS resulted in hypo-responsiveness in both MDM and PSCM, particularly marked for *TNF* expression, and genes that are associated with TNF signaling, including *NFKB1, NAMPT, TNFAIP1, 2* and *8, CXCL2, DUSP2*, and *TNFSF9*. These data confirm that PSCM are LPS responsive and undergo NF-κB-associated tolerance to repeated LPS exposure.

PSCM did not completely mimic the monocyte-derived tolerance program. PSCMs but not MDMs expressed growth factors on LPS restimulation, including *FLT1, PDGFA, VEGFA, CSF1, CSF1R, CSF2, IL32, IL36*, and *TGFB1* (Figure 2 and Supplementary Table 1). Less than 50% of PSCM were recruited to the acute inflammatory response mediated by the MyD88-independent/IRF-dependent pathway, including *IRF1, OAS1, IFIT2* compared to over 80% of MDM (Figure 2D). Review of the pathways differentially expressed by MDM compared to PSCM confirmed that the Reactome IFN signaling pathway scored significantly higher across the MDM time course (Figure 2E). Other targets of intracellular TLR signaling were absent entirely in PSCM, including *IFNB1*, which was inducible in MDM but not PSCM populations. This may have implications for proliferation differences between MDM and PSCM, as CSF1R is targeted by IFNB signaling in monocytes^27^, and further evidences negative feedback of CSF1 signaling in MDM after LPS activation.

Genes that were completely tolerized in the MDM such as *IL1B*, and chemokines such as *CCL3* and *CCL7* were expressed at high levels in acute or resting PSCM, and in the restimulated conditions. These genes were expressed regardless of proliferating status and are predominantly targets of post-transcriptional (AU-response element) regulation, which may be a mechanism that supports acute activation in a macrophage that is also dividing (Figure 2, Supplementary Table 2).

The current model of LPS tolerance is that of an altered chromatin landscape where NF-κB p50 homodimers are stabilized by BCL3, preventing recruitment and binding of NF-κB heterodimers to LPS responsive cytokines^28^. BCL3 nuclear translocation and activation is regulated by proliferation^29^ and is differentially regulated in PCSM but not MDM across the acute phase of the LPS time course, so it is plausible that proliferation of PSCM resets the BCL3-dependant program of LPS tolerance, allowing for ongoing expression of inflammatory mediators in a subpopulation of cells.

## Discussion

In this study we tested two models of cell-intrinsic responses to LPS, identifying proliferation as a driver of macrophage heterogeneity in a minor subset of the MDM, whereas the proliferative subset represented about half of the PSCM population. Endotoxin tolerance in restimulated macrophages took two forms – either in the degree of induction of an inflammatory gene on re-stimulation with LPS or in the proportion of cells capable of expression of an inflammatory mediator. It was the 18 hour-stimulated and restimulated PSCM groups that continued to express cytokines, such as *IL1B*, and chemokines at a high level, possibly because these escaped tolerance in the cells proliferating on first exposure to LPS. TRM have the capacity to self-renew^30^. This homeostatic self-sufficiency of TRM enables the population to be maintained without requiring constant influx of monocytes (Figure 3).

**Figure 3.**
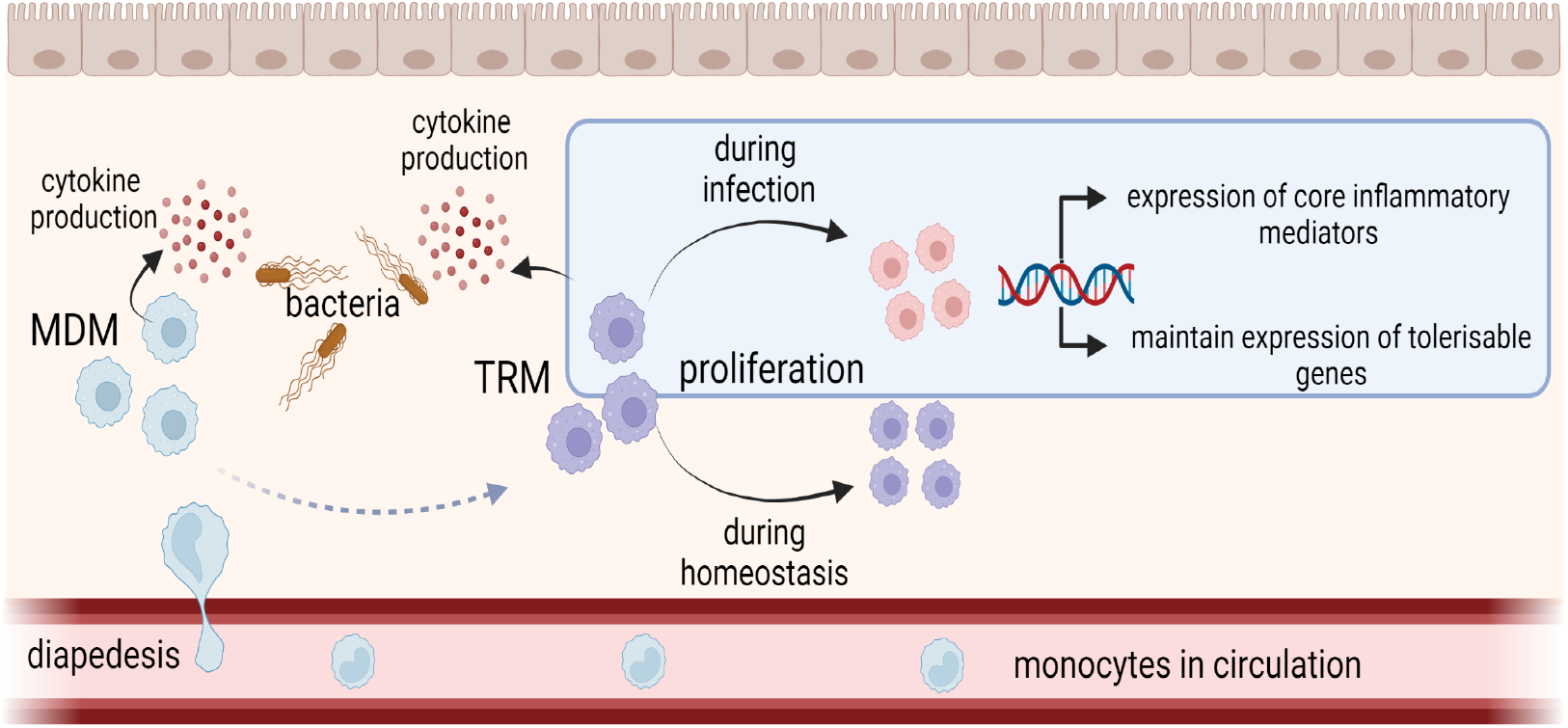
Cartoon of two macrophage populations describing escape from endotoxin tolerance in subpopulations that are proliferating. Created in BioRender.

Prior exposure to a TLR ligand causes hypo-responsiveness of macrophage populations to subsequent stimulations, and LPS/TLR4 drives the largest cohort of tolerized genes^31^. This so-called endotoxin tolerance is regulated by recruitment and stabilization of NF-κB p50 to inflammatory gene promoters, preventing engagement with the transcriptional machinery and silencing the locus^28^. Likewise, prior exposure to low dose pathogenic signals, cytokines, or apoptotic cells can induce ‘trained’ responses in innate immune cells, even days after the initial exposure (reviewed in ^32^). While a key difference between the monocyte-derived MDM and laboratory-derived PSCM is the possibility of prior pathogen exposure in monocytes or precursors^20^, we did not observe any evidence of so-called inflammatory macrophage signatures in the naïve MDM in this study. However, expression of IFN-related pathways in acutely activated MDM is consistent with ‘training’ of MYD88-independent signaling in the MDM rather than PSCM. Activation of IRF-related pathways appears to be part of the monocyte-macrophage transition^33^, but was not recapitulated in the iPSC-derivation protocols.

The experimental design of our macrophage village tested whether LPS tolerance is a consequence of reduced numbers of cells responding to repeated LPS stimulation, or whether NF-κB target genes in the population are uniformly silenced. We observed maintenance of expression of a range of tolerizeable genes in PSCM 18 hour and re-stimulated groups.

Examples of this can be seen in PSCMs through several chemokine and inflammatory genes, including *IL1B, CCL4L2, CCL3* and *IL6* expression, suggesting that in a subset of responsive cells, LPS tolerance had been evaded. Genes previously shown to be stochastically expressed in mouse RAW264.7 cells included *IL6* and *TNF*, with observations that *IL6* was expressed by all RAW264 clones and *TNF* only expressed by 72% of them^13,14^. Similarly, we observed that *TNF* was only expressed in about 75% of human PSCMs, but in 98% of MDMs after acute stimulation with LPS. *IL6* was inducible at very low levels and in few cells in both *in vitro* human macrophage models. We also observed proliferative differences between MDM as a model of recruited and PSCM as a model of resident-tissue macrophages, consistent with the Muldoon model of proliferative-dependent quorum licensing^13^. We hypothesize that the proliferation of resident macrophages ensure that some subsets can evade tolerizing signals to remain responsive to persistent infection and may therefore be the preferred target for persistent inflammation.

## Experimental Procedures

### Cell lines and ethics approvals

Human macrophage derivation was carried out in accordance with The University of Melbourne ethics committee HREC (approvals 1851831, 1646608). Monocytes were isolated from three buffy coats, obtained from the Australian Red Cross Blood Service.

Human pluripotent stem cells (hPSCs) used were: PB001.1 obtained from the Stem Cell Core Facility at the Murdoch Children’s Research Institute (hPSCreg ID: MCRIi001-A ^**34**^); HDF51 (CVCL_UF42) was kindly provided to Dr Andrew Laslett by Prof. Jeanne Loring (The Scripps Research Institute, CA, USA); KOLF2 (Wellcome Trust Sanger Institute; hPSCreg ID: WTSli018-B) cells were kindly provided by Prof. Paul Hertzog (Hudson Institute of Medical Research).

### Pluripotent Stem Cell Differentiation

Human pluripotent stem cells were differentiated into macrophages based on protocol described by ^**18**^. Modifications were as follows: embryoid bodies were kept in rotational cultures without transference to Matrigel plates for adherence, and progenitors collected from week 2 in RPMI-1640 containing L-Glutamine (Life Technologies), 10% Fetal Bovine Serum and 100ng/mL CSF1 for macrophage differentiation (see macrophage differentiation).

### Peripheral Blood Monocyte Isolation

Buffy Coat was diluted with PBS at a 1:3 dilution and underlayed with Histopaque®-1077 (Sigma-Aldrich), centrifuged at 350g for 30 minutes at 24°C with no brake. Peripheral blood mononuclear cells (PBMCs) were isolated from the interphase and washed twice by using MACS buffer with 0.5% heat inactivated Fetal Bovine Serum (FBS) and 2mM EDTA (Invitrogen™ UltraPure™ 0.5M EDTA) and centrifuging at 400g for 5 minutes at 4°C, then resuspended in 40μl MACS buffer per 10^7^ cells. Monocytes were positively selected using Human CD14 MicroBeads (Miltenyi Biotec) and LS Columns, and then used for macrophage differentiation (see below). A cell count and viability were determined using 0.4% Gibco™ Trypan Blue using a hemocytometer.

### Macrophage differentiation

Monocytes or iPSC-derived myeloid progenitor cells were cultured in tissue-culture treated 6-well plates. Cells were cultured in RPMI-1640 medium containing L-Glutamine (Life Technologies) with 10% Fetal Bovine Serum and 100ng/ml recombinant Human M-CSF (R&D Systems) for 4 days.

### Flow Cytometry

Cells were prepared for flow cytometry by first performing a blocking step with 5μl mouse serum (Sigma-Aldrich) then cells were resuspended in FACS Buffer containing 0.5% Bovine Serum Albumin (Sigma-Aldrich) in Hanks Balanced Salt Solution (no calcium, no magnesium, no phenol red) (Thermo Fisher Scientific) and incubated with antibodies (see Table 1). Cell viability was determined using 7-AAD (BioLegend) or PI staining solution (Invitrogen eBioscience). Gating was performed based on Fluorescence Minus One. Isotypes were included as a control (Table 1). Compensation was carried out by generating a matrix using compensation beads (BD™ CompBead Plus; Anti-79 mouse Ig, κ). Cell analysis was carried out on a CytoFLEX S flow cytometer (Beckman Coulter) using CytExpert acquisition software. Post-acquisition analysis was performed with FCS Express 7 flow cytometry software.

**Table 1:** Flow Cytometry Antibodies

| Table 1 : Flow Cytometry Antibodies |  |  |
| --- | --- | --- |
| Marker | Antibody | Isotype control |
| CD14 | Brilliant Violet 421™ anti-human CD14 Antibody<br>(BioLegend; Cat. No. 325628) | Brilliant Violet 421™ Mouse IgG1, κ Isotype Ctrl Antibody<br>(BioLegend: Cat. No. 400157) |
| CD16 | Brilliant Violet 605™ anti-human CD16 Antibody<br>(BioLegend: Cat. No. 302040) | Brilliant Violet 605™ Mouse IgG1, κ Isotype Ctrl Antibody<br>(BioLegend: Cat. No. 400161) |
| CD45 | APC anti-human CD45 Antibody<br>(BioLegend: Cat. No. 368512) | APC Mouse IgG1, κ Isotype Ctrl (FC) Antibody<br>(BioLegend: Cat. No. 400121) |
| CD11b | Brilliant Violet 711™ anti-human CD11b Antibody<br>(BioLegend: Cat. No. 301344) | Brilliant Violet 711™ Mouse IgG1, κ Isotype Ctrl Antibody<br>(BioLegend: Cat. No. 400167) |
| HLA-DR | Alexa Fluor® 488 anti-human HLA-DR Antibody<br>(BioLegend: Cat. No. 307619) | Alexa Fluor® 488 Mouse IgG2a, κ Isotype Ctrl Antibody<br>(BioLegend: Cat. No. 400233) |

### Stimulation assay

100,000 cells of CD14+ HDF51-, PB001.1- and KOLF2-derived progenitors, or primary CD14+ monocytes from 3 different donors, were pooled then seeded into 6-well dishes in Gibco™ RPMI-1640 media (Thermo Fisher Scientific) with 10% heat inactivated FBS (Thermo Fisher Scientific) and 100ng/ml M-CSF (R&D Systems). On day 4, media was changed, and cells left to settle for 2 hours before stimulating with 10ng/mL LPS. After 18 hours, tolerance wells were washed twice with Gibco™ PBS (Ca2+Mg2+ free; Thermo Fisher Scientific) before fresh media was added and cells left to settle for 2 hours and re-stimulated with 10ng/mL LPS for 2 hours. Cells were washed with Gibco™ PBS (Ca2+Mg2+ free; Thermo Fisher Scientific) before incubating with macrophage cell release for 5 minutes at 37°C and collected in media, centrifuged at 350g for 5 minutes and resuspended and put through a Flowmi™ strainer (before cell counts were carried out for sample preparation, see 10x Illumina Single Cell Library Preparation and RNA Sequencing).

### 10x Illumina Single Cell Library Preparation and RNA Sequencing

Single cell RNA-sequencing libraries were prepared using the Chromium Next GEM Single Cell 5’ v2 (Dual Index) protocol as per manufacturer’s instructions. Cell suspensions were prepared for a target recovery of 6000 cells per sample. Libraries were quality checked and quantified using the Agilent Tapestation D5000 High Sensitivity Tape. Two sequencing runs were performed using Novaseq6000 at University of Melbourne Centre for Cancer Research. Data processing and merging of sequencing runs were carried out through CellRanger v6.0.2 pipeline to generate files for analysis. Downstream analysis was carried out using Seurat package v4.0.5 ^35^.

### Donor Deconvolution

The raw number of reads that were associated with each barcode were counted, and only cells with more than 2000 reads associated were kept for further analysis. We applied cellSNP-lite (with options: --minMAF 0.1, --minCOUNT 5)^36^ for getting the allele counts for the list of common SNPs in the population identified from the 1000 Genome Project^37^. We subsequently applied Vireo^38^ for demultiplexing the cell barcodes into 3 donors or cell lines used for each sample. The same donors were then matched by their genotype similarity (Supplementary Figure 1).

### Differential gene expression analysis

A pseudo-bulk approach was used to identify differentially expressed genes^39^. The gene counts in cells for each donor were aggregated to form bulk samples, and we applied differential expression analysis using methods designed for bulk RNA-seq^40,41^. Briefly, 24 bulk samples were created after summing gene counts for 3 donors in each time point, and lowly expressed genes were filtered out. The gene counts for all samples were then normalized using TMM (trimmed mean of M-values) ^40^. Differential genes were identified by contrasting samples between sample conditions, and the final list of DE genes in each contrasting group were obtained by the “treat” method with log(2) fold change > 1^42^.

### Gene Set and Pathway Enrichment Analysis and ARE Motif Analysis

AU-rich element motifs were evaluated using the ARE motif database ^43^. The search space was restricted to 3’UTR motifs. Gene Set Enrichment Analysis (GSEA) was performed using the MSigDB Hallmark gene sets ^44^. Pathway enrichment used InnateDB ^45^ or Reactome ^46^ on upregulated genes ≥1.5 Log2FC and p.value ≤0.05. Interferon module scores were calculated with the Seurat function AddModuleScore using the Reactome interferon signaling pathway gene set, obtained from the msigdbr package. Bar plots showing the average expression of example genes were derived from the output of the Seurat DotPlot, taking the percentage of cells per group expressing each gene of interest (bar height) and the unscaled average expression per group (colour).

## Author Contributions

N.R. experimental design, data generation and analysis, writing

L.J.G. data analysis

R. L. SNP deconvolution and analysis

Y.D.J.P library preparation, data processing

P.W.A. data analysis

S. M.G experimental resources

A.L.L. experimental resources

D.M. experimental resources

C.A.W. experimental design, data analysis, writing, project funding

## Acknowledgements

The authors thank the Genomics Platform Group at The University of Melbourne Centre for Cancer Research (UMCCR) sequencing core for sequencing of single-cell libraries, and Prof. Oliver Hofmann for computational workflows and project discussions. The authors thank Dr Catherine King (10x Genomics) and Dr Paul Gooding (Millenium Science) for their training and assistance in preparation of 10x libraries. The authors thank Dr Jarny Choi and Dr Michael Clark, University of Melbourne, for reading and commenting on the manuscript.

## Data and code availability

Single cell sequencing data are available through (GSE210175). Code used for analysis is available at: https://github.com/wellslab/invitrosinglecell/blob/main/Analysiscode.R

---

**Supplementary Figure 1:**
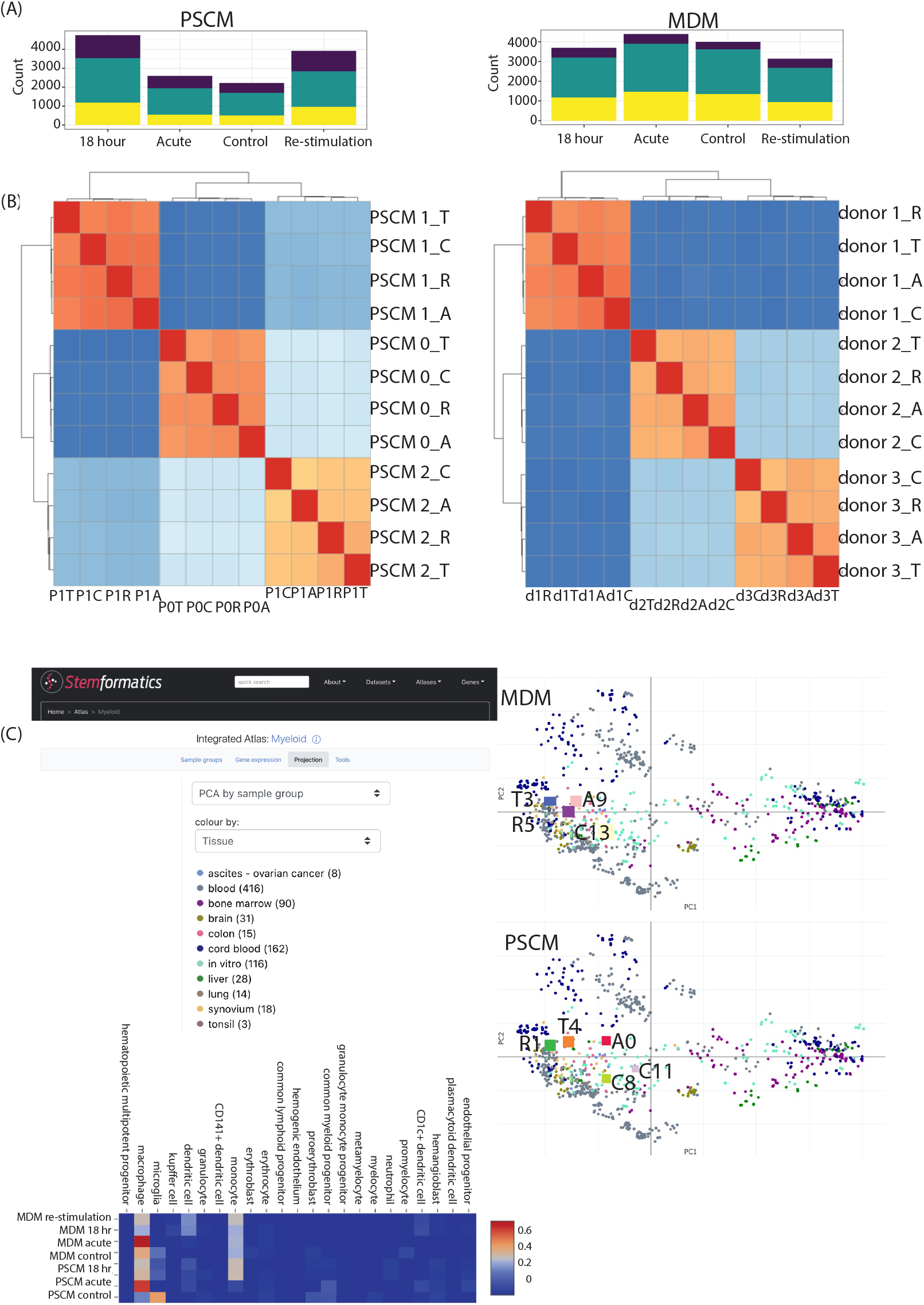
Quality assessment of RNAseq libraries. A: donor contributions to PSCM or MDM sequencing libraries. B: Correlation of sequencing tags labelled by experimental condition (barcode) or donor ID. C: Screenshot of projection of PSCM and MDM libraries onto the Stemformatics Myeloid atlas, and Cabyara analysis of most similar reference atlas samples.

## Supplementary Material

**Supplementary Table 1: Differential expression analysis of MDM and PSCM groups. Accompanies Figures 1 and 2.**

Average and differential expression analysis between MDM and PSCM groups. Average expression, Log2FC, p value, and adjusted p value.

**Supplementary Table 2: Average expression of TLR4 signaling components. Accompanies Figure 2A.**

Average expression of components of TLR4 signaling pathway. Average expression shown for all identified seurat clusters.

